# Standard-flow targeted proteomics for quantification of enzymes in glucosinolate biosynthetic pathways

**DOI:** 10.1101/2024.11.15.623753

**Authors:** Daniel Vik

## Abstract

Insights gained from quantitative proteomics studies show that the relationship between transcript and protein abundance is not generally linear. Quantitative measurement of both proteins and transcripts is required to reveal their actual relationship. Whereas quantitative analysis of transcripts is widely established, there is a need for easily accessible protein level technologies. Mass spectrometry-based proteomics is a promising solution, provided that it can be made accessible to a broader user group than it currently is. Developments in proteomics methodology show that relatively simple instrumentation allows for targeted proteomics which provides a reliable and accessible technology platform for protein quantification. I here describe the successful implementation of standard-flow targeted proteomics and its use to quantify 12 enzymes in the glucosinolate biosynthetic pathways in Arabidopsis thaliana. Enabled by targeted proteomics, I analyzed a series of transcription factor mutants and found that transcript and protein generally correlated well under these unchallenged conditions. When I treated wild-type plants with the phytohormone methyl-jasmonate, I observed uncoupling of metabolite, transcript and protein levels, suggesting post-transcriptional and/or -translational in addition to transcriptional regulation.

## 1 Introduction

Mass spectrometry (MS)-based proteomics is a powerful approach to study proteins. Conventional shotgun proteomics also known as data-dependent acquisition (DDA) is among the most widely used proteomics approaches. Shotgun proteomics typically identifies thousands of proteins in a single sample that are quantifiable by various means (Ong and Mann, 2005). DDA experiments, however, require highly specialized equipment, which can be difficult to access and demanding to maintain. In addition, DDA experiments generally suffer from low repeatability and reproducibility (Bell et al., 2009; Tabb et al., 2010). Another critical limitation arises from the huge differences in abundance between proteins in biological samples, which makes it challenging to identify lowly abundant proteins (Zubarev, 2013). This problem is especially prevalent when working with samples were the examined analytes span many orders of magnitude, such as plasma samples, – where hemoglobins and serum albumins dominate the proteome (Anderson, 2002), – or as in the case of most plant samples, which are challenging due to massive amounts of photosynthetic proteins (Smaczniak et al., 2012; Gupta et al., 2015). An alternative proteomics method, selected reaction monitoring (SRM) assays – or “targeted proteomics” – has gained popularity in recent years, as it alleviates some of the problems associated with DDA experiments (Doerr, 2010). These targeted peptide assays exploit the mass-filtering capabilities of triple quadrupole MS instruments. This allows for highly selective analysis and enables detection and quantification of a specific subset of the proteome, without extensive sample fractionation or enrichment (Anderson and Hunter, 2006; Wienkoop and Weckwerth, 2006; Picotti et al., 2009; Ito et al., 2011; Batth et al., 2014; Ito et al., 2014; Taylor et al., 2014). Stable isotope-labeled peptides are used as internal standards and their relative intensities form the basis for quantification which is greatly facilitated by the developments in chromatogram-centric data analysis software (MacLean et al., 2010). An added benefit of targeted proteomics is its compatibility with standard-flow liquid chromatography (LC) (compared to challenging, high sensitivity nano-flow LC), which provides increased robustness with shorter gradients (Domanski et al., 2012; Percy et al., 2012; Batth et al., 2014; González Fernández-Niño et al., 2015). Standard-flow-driven targeted proteomics thus offers several benefits: i) stable and fast chromatography, ii) high selectivity and iii) simple data analysis. In addition, standard-flow LC instruments coupled to triple quadrupole MS detectors are widespread in many labs that carry out metabolic analyses of plants, and thus represent immense potential for a broader application of hypothesis-driven, targeted proteomics in plant biology.

Transcript analyses are among the preferred methodologies to study the regulation of biosynthetic pathways, and co-expression analysis (i.e. the guilt-by-association principle) has been quintessential for elucidating biochemical pathways (Hirai et al., 2005; Sønderby et al., 2010b; Celedon and Bohlmann, 2016; Morris et al., 2016). An example are the glucosinolate (GLS) defense compounds that are characteristic of brassicaceous plants, including the model plant Arabidopsis thaliana (hereafter Arabidopsis). Their biosynthetic pathways have been studied extensively via transcript analysis, which facilitated the identification of critical enzymes and regulators in Arabidopsis (Hirai et al., 2007; Hansen et al., 2008; Sønderby et al., 2010a; Sønderby et al., 2010b; Schweizer et al., 2013; Burow et al., 2014; Frerigmann and Gigolashvili, 2014). An assumption underlying co-expression analyses is that changes in transcript levels represent similar changes in enzyme abundance. This assumption is being challenged by proteomics studies, showing that transcript and protein generally show weak correlation (Gygi et al., 1999; Orntoft, 2002; Maier et al., 2009; Baerenfaller et al., 2012). Recent studies have examined the GLS pathway by comparing metabolomics and proteomics data (He et al., 2013; Mostafa et al., 2016), but the relationship between transcript and enzyme remains elusive. Understanding the relationship between transcript and protein is an important aspect in understanding pathway orchestration. More importantly, insights into the transcript-protein relationship have implications for our interpretation of previous and future data, and the capacity of transcript levels to reflect gene expression and function.

Enabled by standard-flow-driven targeted proteomics, I here report the comparison of transcript and enzyme levels from the two GLS core pathways in Arabidopsis that convert tryptophan and chain-elongated methionine into the corresponding indole and aliphatic GLS, respectively (Halkier and Gershenzon, 2006; Sønderby et al., 2010b). A substantial number of peptide assays are required to cover the core pathways, which are composed of 12 enzymes. In addition, I include peptides covering the CYP71B15/PAD3 enzyme – the last enzyme in the biosynthesis of the phytoalexin camalexin (Schuhegger et al., 2007; Bottcher et al., 2009). I analyzed a series of GLS biosynthetic mutants lacking bHLH and R2R3 MYB transcription factors showing strongly reduced transcript levels of GLS biosynthetic genes (Hirai et al., 2007; Sønderby et al., 2007; Schweizer et al., 2013; Frerigmann and Gigolashvili, 2014). The myb mutants were previously reported to show discrepancies between transcript levels and the corresponding GLS output which suggests alternative regulatory mechanisms at the post-transcriptional and/or post-translational level (Sønderby et al., 2010a; Frerigmann and Gigolashvili, 2014). In addition, I investigated the dynamic behavior of transcripts, proteins and metabolites in response to external application of methyl-jasmonate (MeJa) to identify the regulatory levels responsible for induced accumulation of GLS defense compounds. Our study further provides a case study of how to establish a targeted proteomics workflow in a laboratory equipped for metabolite analysis by standard-flow LCMS.

## 2 Results

### 2.1 Optimization of settings for targeted proteomics

A singular set of parameters for chromatography and ionization that is optimal for all peptides does not exist. Instead, the challenge is to find a set of parameters that broadly enables sufficient ionization of all analytes, without compromising any individual analyte, referred to as the Pareto-optimal solution. Using a set of commercially available synthetic peptides, I examined the different parameter settings (figure 1). Initially, I tested the voltage for electrospray ionization between 1500 V and 4800 V (figure 1 A). As no single voltage was optimal for all peptides, I selected an intermediate voltage of 3200 V. I then examined the effects of the flow rate. Optimization of this factor has been reported to allow standard-flow LC to achieve sensitivity similar to nano-flow LC, which is crucial for quantification of low-abundant proteins (Percy et al., 2012). Our results show that flow rates above 300 µL/min improve peak shape, as seen clearly for the ADEGISFR peptide (figure 1B). Beyond the initial improvement achieved by increasing flow rates from 300 µL/min to 400 µL/min I did not observe any obvious benefits from increased flow rates. Further examination of the settings for the ion source included nebulizer gas flow, probe gas flow and probe temperature. Comparing the effect of nebulizer gas flow between three peptides, it is evident that higher nebulizer gas flow yields larger peak areas; almost two-fold increase in signal when increasing flow from 30 to 60 psi (figure 1C). Much less pronounced are changes of the probe gas flow, which show little difference in peak areas (figure 1D). Higher probe temperatures generally lead to larger peak areas – peaking around 325 °C (figure 1C). The above systems settings present the Pareto-optimal settings for all tested synthetic peptides on the present instrumentation. Based on these initial tests, I assume that these settings can be applied to a broader range of peptides with similar results.

**Figure 1.**
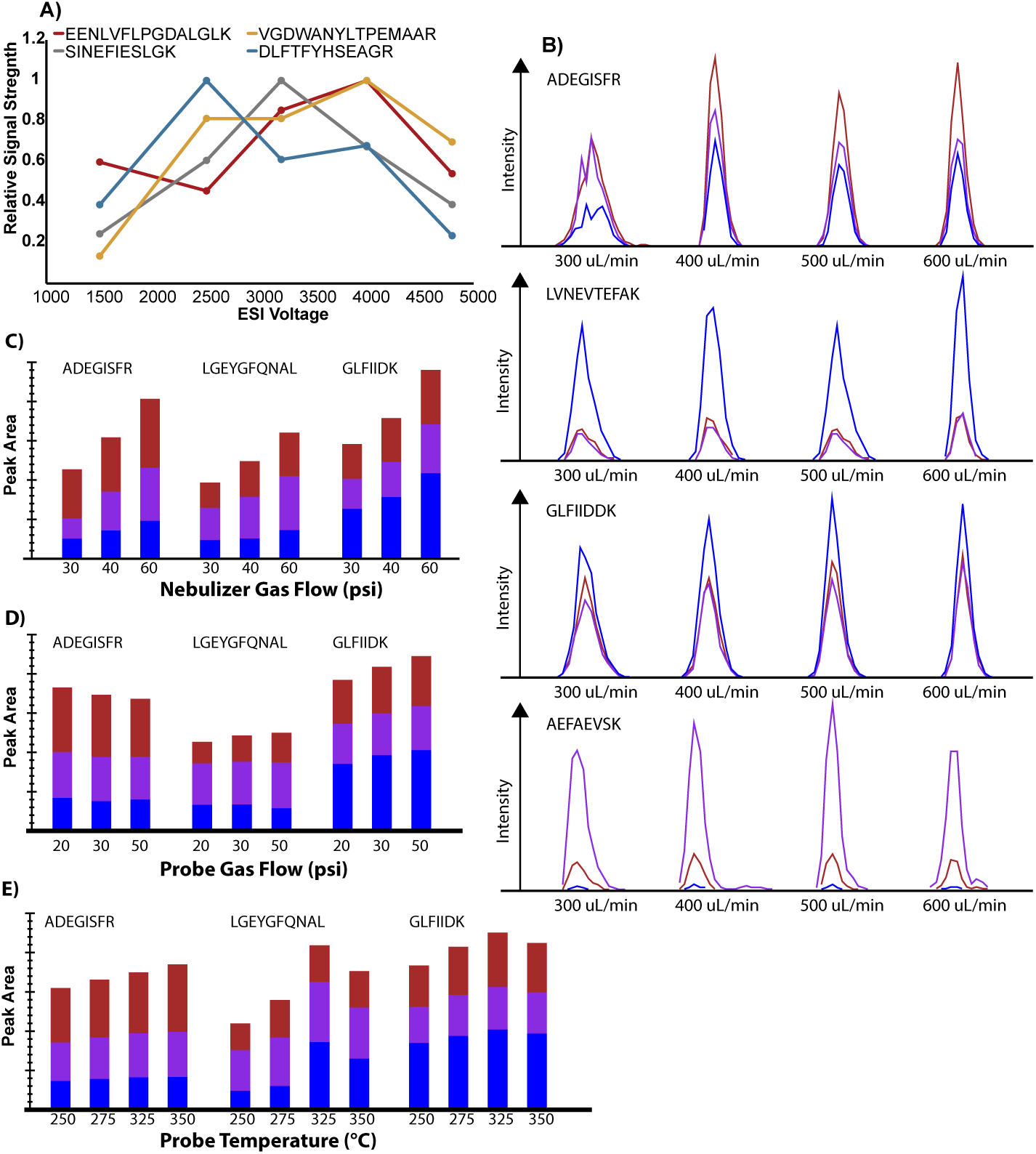
Examination of optimal ionization settings for selected synthetic peptides on a triple quadrupole mass spectrometer. A) Survey of electrospray ionization voltage. B) Effect of flowrate on peak shape and intensity. C) Nebulizer gas flow. D) Probe gas flow. E) Probe temperature. Representative synthetic peptides are shown; the colors correspond to three different transitions for each peptide (B-E).

### 2.2 Design of SRM assays for GLS biosynthetic enzymes

A critical step in designing SRM assays for quantification of peptides is the choice of proteotypic peptides, i.e. peptides whose sequence unequivocally represents a distinct protein. The quality of the assay depends entirely on the nature of these peptides. Searching through published DDA proteomics data is an effective way to find good candidate peptides, but these datasets are not complete, and many proteins are not represented, particularly in the case of low-abundant proteins. If no peptides have been reported, in silico tryptic digest combined with the various prediction algorithms such as CONSeQuence, the Arabidopsis Proteotypic Predictor server and ExPASy can help select candidate peptides (Wilkins et al., 1997; Eyers et al., 2011; Taylor et al., 2014). For the SRM assays of enzymes in the GLS and CYP71F15/PAD3 in the camalexin pathway, I utilized the pep2pro database (Baerenfaller et al., 2011; Hirsch-Hoffmann et al., 2012), which covers many DDA experiments and more than 27400 loci of the TAIR10 Arabidopsis genome assembly. For several of our proteins of interest, peptides had been identified in DDA experiments with Arabidopsis. However, CYP79B2, CYP79B3, CYP79F1, CYP79F2 and CYP71B15/PAD3 had almost no reported peptides. These proteins were thus subjected to in silico trypsin digest and the resulting proteotypic peptides were ranked according to their predicted ionizability using the Arabidopsis Proteotypic Predictor server (Taylor et al., 2014). The highest scoring peptides were synthesized with an N-terminal isotope-labeled lysine or arginine, and were subjected to collision energy optimization via direct infusion of the synthetic peptides into the triple quadrupole MS. The transitions – i.e. pairs of specific m/z values for a parent peptide and one of its fragment ions – were either picked based on available spectral libraries such as the Global Proteome Machine (Craig et al., 2004) or individual experiments found in pep2pro. If no spectra were available, I selected the 5-7 fragment ions with highest m/z values based on the primary sequence of the respective proteotypic peptide. I screened a total of 88 peptides, covering the core structure biosynthesis of both tryptophan- and methionine-derived GLS as well as camalexin biosynthesis, the latter being represented by peptides of CYP79B2, CYP79B3 and CYP71B15/PAD3 (Bottcher et al., 2009; Sønderby et al., 2010b; Jensen et al., 2014). Peptides were selected based on the following screening criteria: i) good signal for at least three transitions via direct infusion into the triple quadrupole MS; ii) symmetrical peak shape and iii) three transitions with no interferences from the Arabidopsis proteome determined by manual peak inspection.

Our screening using synthetic peptides yielded 30 functional SRM assays (table 1). All assays developed with synthetic peptides also detected endogenous peptides from Arabidopsis samples, except for four cytochrome P450s: CYP79B2, CYP79B3, CYP79F2 and CYP71B15/PAD3. However, these assays were kept in the method as the synthetic peptides gave good signal, and our inability to detect these endogenous peptides could have been due to low basal levels of the respective proteins. Except for the peptides FASIVPVLTLAGISK (SUR1), DAITPGSYFGNEIPDSIAIIK (GGP1), VVNLCSFETLK (SOT18) and TMGTSSYGEHFMK (CYP79F2); all peptide assays showed a lower limit of quantification around or below 5 fmol of injected peptides in samples composed of Arabidopsis protein extracts representative of those used throughout all subsequent analyses (figure S1).

**Table 1.**
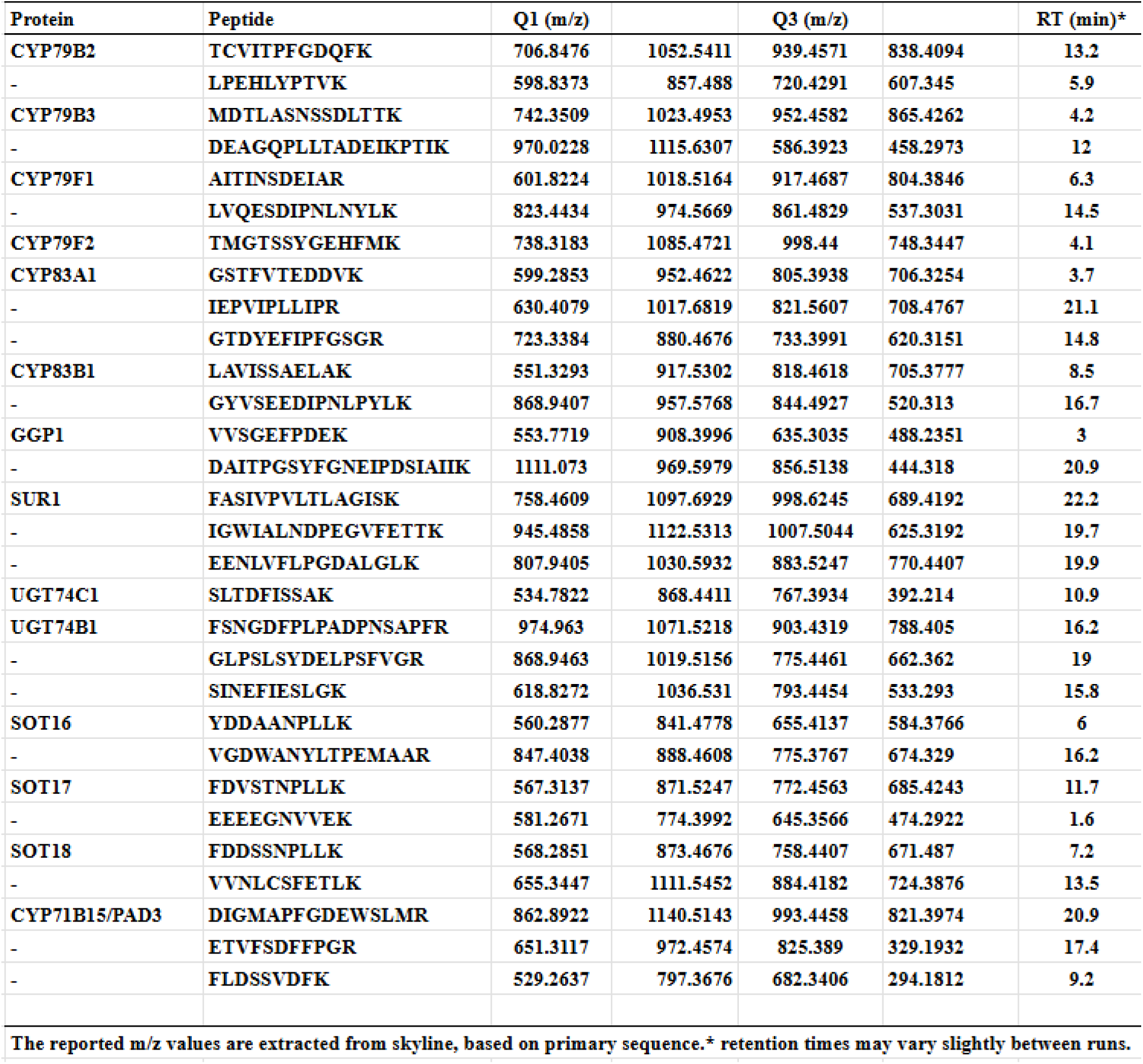
SRM assays for GLS and camalexin biosynthetic enzymes.

### 2.3 Considerations for quantification of enzymes

Peptide quantification is based on the ratios between endogenous and synthetic peptides that are being spiked in in a consistent amount across samples. A basic assumption for inference of protein abundance is that the abundance of a peptide corresponds to that of the parent protein; however, this is not necessarily the case. Both technical (e.g. trypsin digest efficiency or differential peptide adsorption to materials) and biological factors (e.g. post-translational processing or alternative splicing) can distort the relation between protein and peptide abundance. High confidence quantification can be obtained by measuring two or more peptides of a protein. When the signals obtained from two peptides are perfectly correlated across sample replicates, the protein abundance can be confidently inferred from any of the two peptides as seen for CYP79F1 (figure 2A). However, occasionally two peptides deviate from each other, and it can be challenging to justify inference by either peptide. This is seen for the peptides of SOT16 (figure 2B). For SOT16, the peptide-peptide correlation is weak, which makes high confidence quantification using any of these two peptides difficult (figure 2B). R-squared values for the best correlated peptide pairs for each protein can be found in table 2 and corresponding peptide-peptide plots are shown in supplemental figure S2. Besides the peptides showing low degree of correlation, care must be taken when quantifying CYP79F2 and UGT74C1, as only a single peptide for each protein met our screening criteria making it impossible to confidently infer protein abundance. In cases where a correlating peptide pair is unavailable as in the case of SOT16, inference of protein abundance may still be meaningful, however, care must be taken to interpret these data only as suggestive of protein abundance.

**Table 2.**
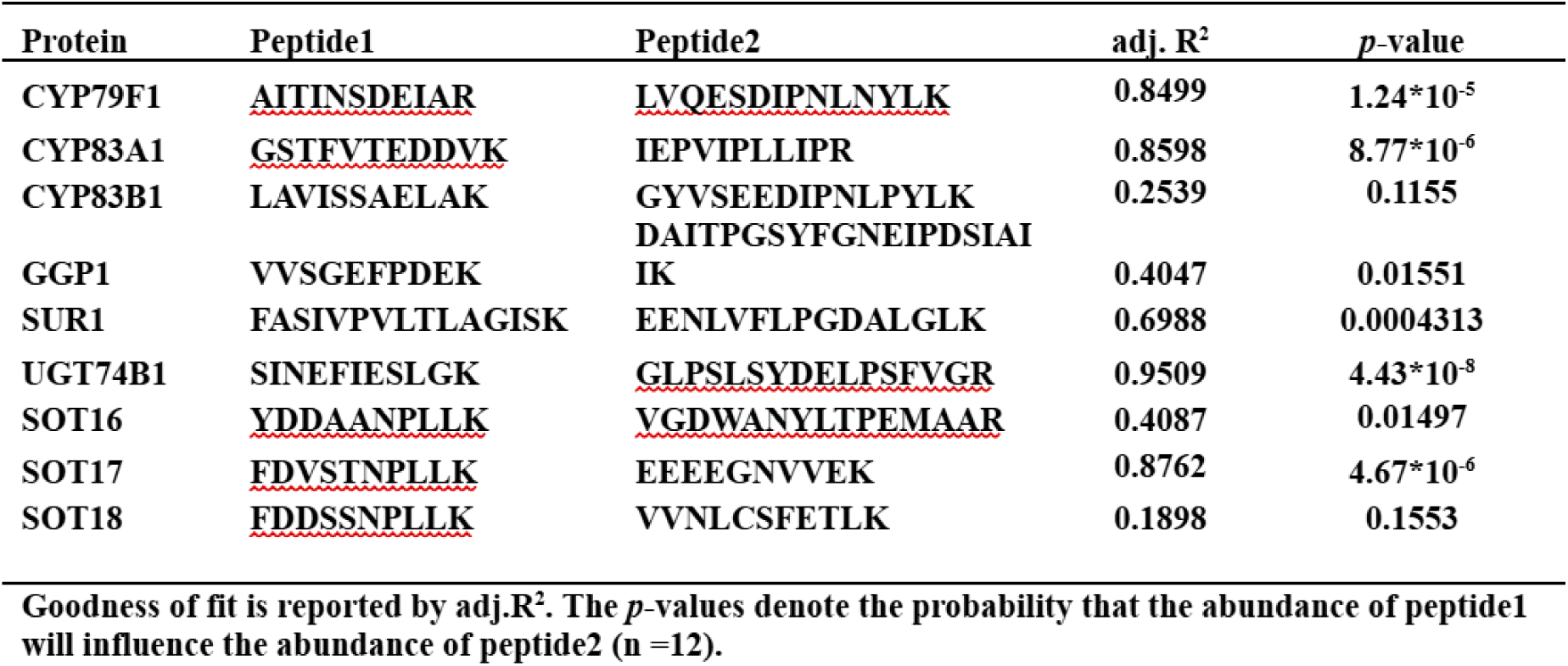
Correlation between peptides from the same protein. Pairs of peptides were quantified and fitted to a linear model.

**Figure 2.**
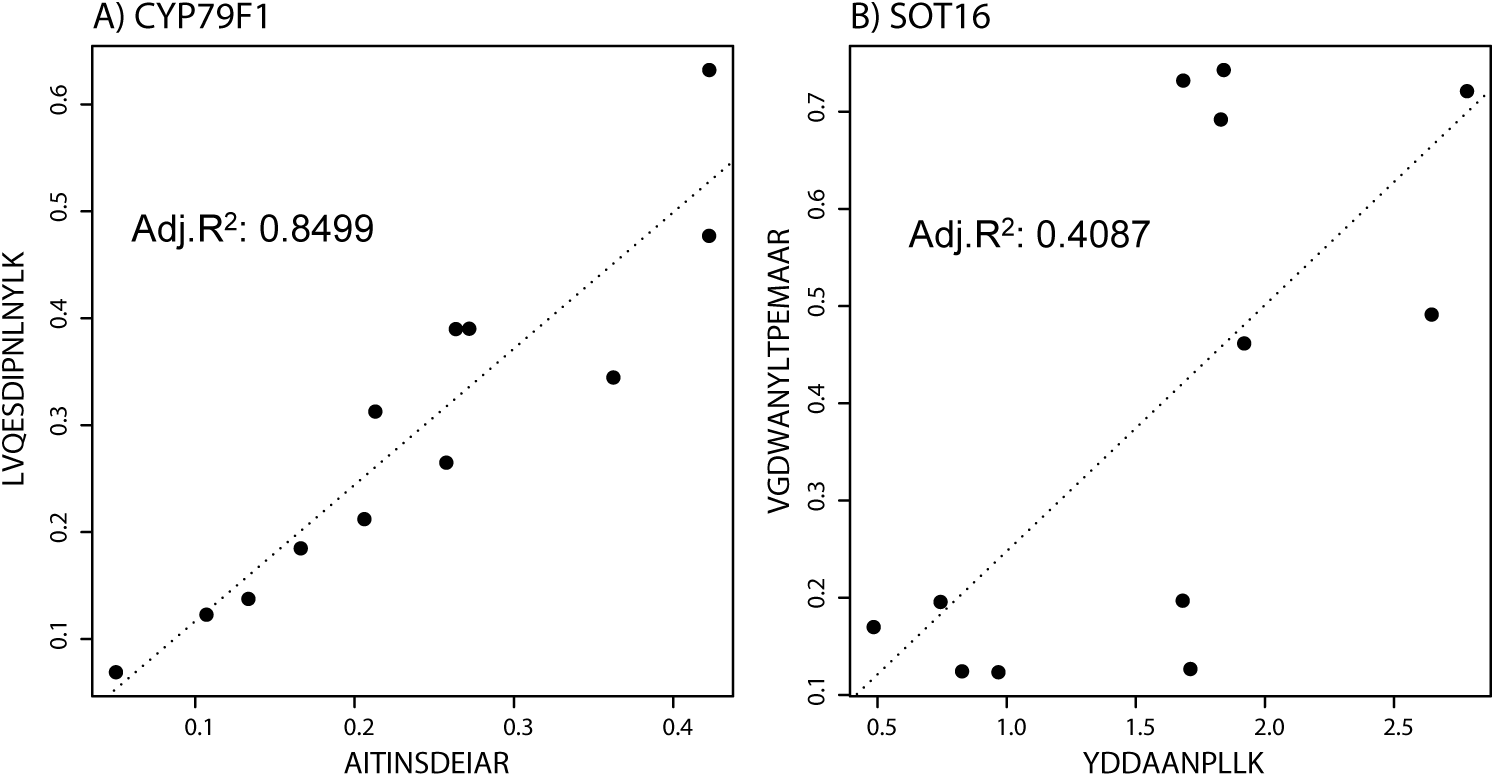
Correlation of peptides from the same protein for confident inference of abundance. The relative abundance of two quantified peptides is plotted against each other. The dashed lines represent linear regressions from which R^2^ was determined (table 2). A) CYP79F1 and B) SOT16. Additional peptide-peptide plots are shown in supplemental figure S2.

### 2.4 Comparison of protein and transcript levels in aliphatic- and indole GLS-specific transcription factor mutants

Towards our goal of characterizing the GLS biosynthetic pathways at the enzyme level and examining the correlation between transcript and protein level, I analyzed a set of mutant lines whose GLS phenotypes have previously been documented, both at metabolite and transcript level. The myb28/29 double mutant accumulates only traces of aliphatic GLS (AG), and slightly increased levels of indole GLS (IG) (Sønderby et al., 2007; Sønderby et al., 2010a). Similarly, only trace levels of IG are left in the myb34/51/122 triple mutant, whereas the level of AG is unaffected or slightly increased (Frerigmann and Gigolashvili, 2014). The myc2/3/4 triple mutant is completely devoid of any GLS (Schweizer et al., 2013). Using SRM assays to quantify peptides of the biosynthetic enzymes, I found that the cytochrome P450s representing the first two steps in the pathways are severely affected in the respective mutants, with levels close to our detection limit (figure 3).

**Figure 3.**
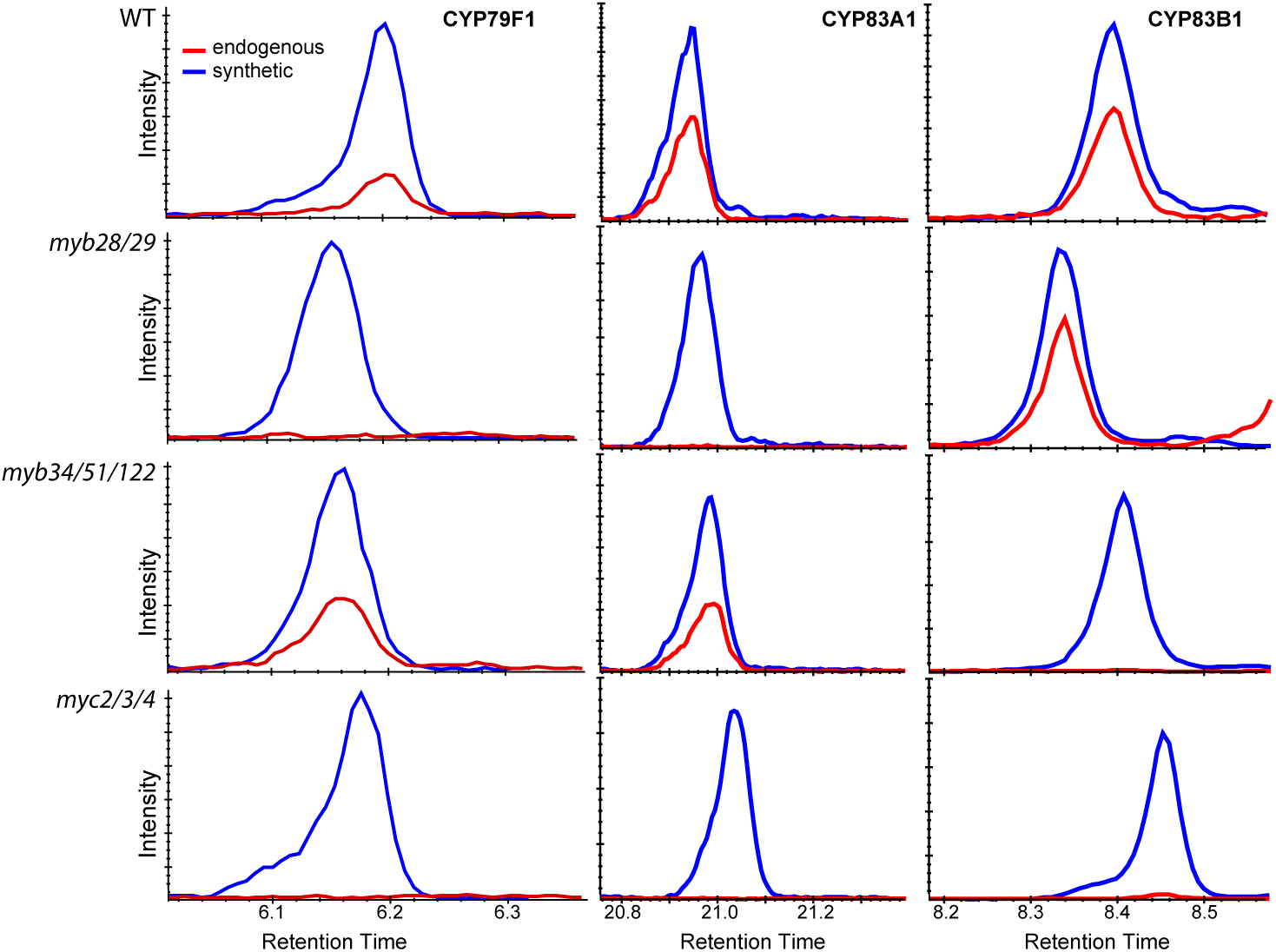
Chromatograms from SRM analysis of GLS-associated cytochrome P450s in transcription factor mutants. Left column shows chromatograms of the AITINSDEIAR peptide representing CYP79F1. Middle column shows IEPVIPLLIPR representing CYP83A1. Right column shows LAVISSAELAK representing CYP83B1. The four rows represent the genotypes analyzed (from the top row down: WT, *myb28/29*, *myb34/51/122* and *myc2/3/4*). Endogenous peptides are shown in red, and the isotope-labeled, synthetic peptide standards are shown in blue. The y-axis is normalized to labeled peptide standards.

Taking advantage of solid-phase extraction of RNA, I purified both RNA and protein from the same samples, and performed SRM assays for protein abundance and quantitative real-time PCR (qRT-PCR) for transcript abundance. This enabled us to compare the relative changes occurring in the different transcription factor mutants (figure 4). Expression of the GLS-associated cytochrome P450 enzymes was severely affected and the protein levels followed transcript abundance closely. Interestingly, CYP83B1 – an enzyme in IG biosynthesis – shows increased levels of transcript and protein in the myb28/29 mutant relative to WT (figure 4C, table 3 and table 4).

**Table 3.** Statistical analysis of transcript levels in mutants. Comparison of transcript levels in three different mutant lines relative to WT. Table contains p-values from *posthoc* test (Welch two sample t-test, n = 12).

| Gene | *myb28/29* |  | *myb34/51/122* |  | *myc2/3/4* |  |
| --- | --- | --- | --- | --- | --- | --- |
| *CYP79F1* | 2.0*10<sup>-5</sup> | *** | 0.7006 |  | 2.5*10<sup>-5</sup> | *** |
| *CYP83A1* | 2.2*10<sup>-7</sup> | *** | 0.9942 |  | 2.0*10<sup>-7</sup> | *** |
| *CYP83B1* | 0.0186 | * | 3.2*10<sup>-6</sup> | *** | 8.9*10<sup>-5</sup> | *** |
| *GGP1* | 0.9936 |  | 0.0138 | * | 0.0005 | *** |
| *SUR1* | 0.4657 |  | 0.0141 | * | 0.0222 | * |
| *UGT74C1* | 0.0093 | ** | 0.0101 | * | 0.0029 | ** |
| *UGT74B1* | 0.7712 |  | 0.0002 | *** | 3.0*10<sup>-5</sup> | *** |

\**p*-value < 0.05, \*\**p*-value < 0.01, \*\*\**p*-value < 0.001. The associated ANOVA-table can be found in supplemental table S4.

**Table 4.**
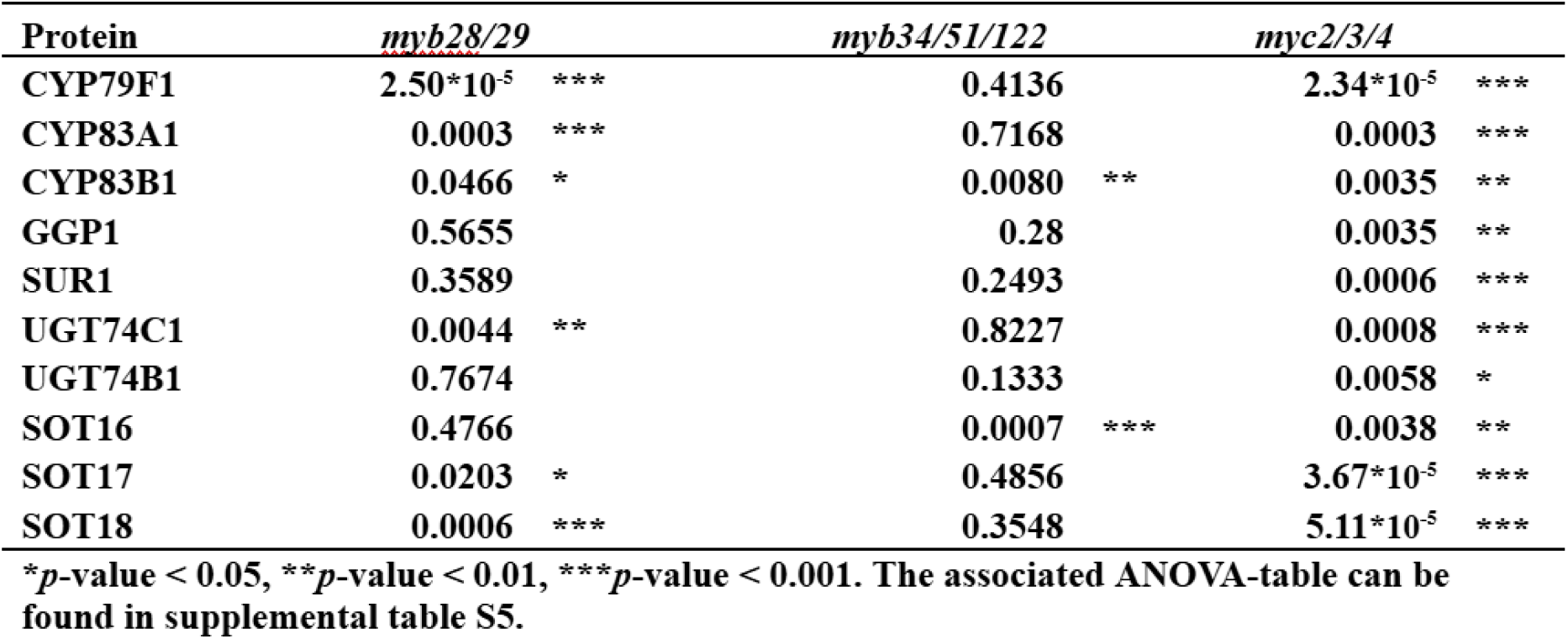
Statistical analysis of protein levels in mutants. Comparison protein levels in three different mutant lines relative to WT. Table contains p-values from *posthoc* test (Welch two sample t-test, n = 12).

**Figure 4.**
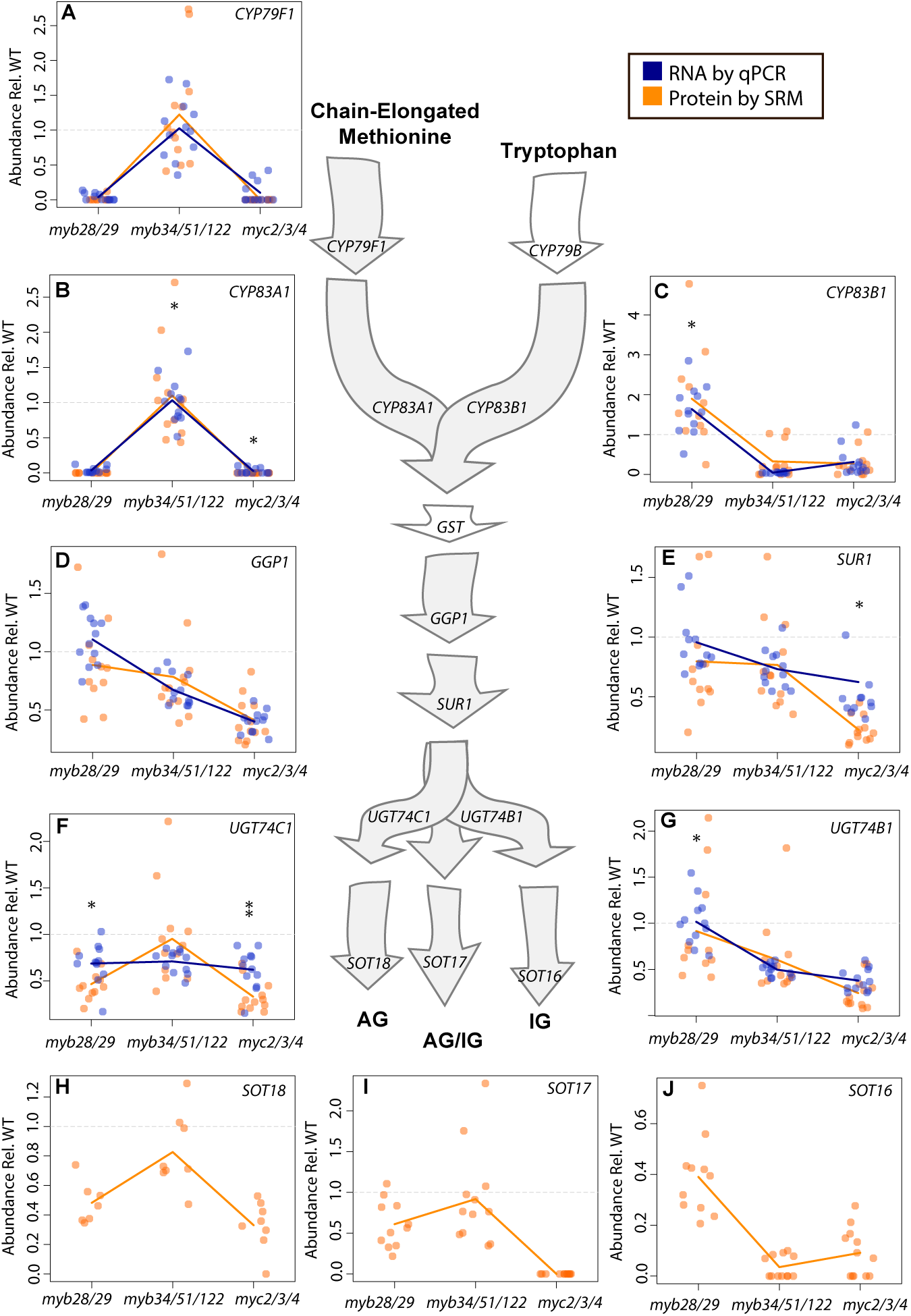
Levels of GLS biosynthetic enzymes and transcripts in different biosynthetic mutants. Schematic representation of the core metabolic pathways of AG (left) and IG biosynthesis (right). WT-normalized, relative abundance of protein (yellow) and transcript (blue) is shown for the three analyzed mutants (*myb28/29*, *myb34/51/122* and *myc2/3/4*). Protein abundance is inferred from individual peptides seen in Table S5. Dots represent biological replicates (n = 12; three experimental rounds each with four biological replicates); lines represent the mean. The grey dashed line represent the WT level set to 1. Asterisks denote statistical significant differences between transcript and protein levels for the individual genotypes (\**p* -value < 0.05, \*\**p* -value < 0.01, ANOVA and Welch two sample t-test as *posthoc* test, supplemental table S3).

Several of the enzymes in our assays are shared between AG and IG biosynthesis, including GGP1 and SUR1. Both transcript and protein levels of GGP1 and SUR1 showed a severe decrease in the myc2/3/4 mutant, but more modest changes in transcript in the myb34/51/122 mutant (figure 4D, E, table 3). GGP1 displays good correlation between transcript and protein (figure 4D and supplemental table S3), but for SUR1 I find a disproportionate reduction in protein in the myc2/3/4 mutant, compared to the corresponding transcript (figure 4E, supplemental table S3). The immediate step following SUR1 is catalyzed by either UGT74B1 or UGT74C1. UGT74B1 shows protein and transcript profiles that resemble those of the shared enzymes GGP1 and SUR1. It is not reduced in the AG mutant myb28/29, but shows a significant transcript reduction in the IG mutant myb34/51/122, and reduction in both transcript and protein levels in the myc2/3/4 mutant (figure 4G, table 3 and table 4). For UGT74C1, I found that the abundance of the protein is not reduced in myb34/51/122 but significantly reduced in myb28/29 and myc2/3/4 (table 4). UGT74C1 transcripts on the other hand, are significantly reduced in all mutants – with no difference between mutants (figure 4F and table 3).

The last step in GLS core structure biosynthesis is the addition of a sulfate group catalyzed by three paralog sulfotransferases: SOT16 shows a preference for intermediates in IG biosynthesis, SOT17 is shared between both pathways and SOT18 shows a preference to AG precursors (Piotrowski et al., 2004; Klein et al., 2006). Their proposed functional specificity is reflected by the expression profiles of the proteins. Assuming that the peptide VGDWANYLTPEMAAR reflects SOT16 protein levels, SOT16 is reduced in all mutants except myb28/29 (figure 4J and table 4). SOT18 is reduced in myc2/3/4 and myb28/29, but unaffected in the myb34/51/122 triple mutant (figure 4H and table 4). SOT17 is below detection limit in myc2/3/4, and significantly reduced in myb28/29 and myb34/51/122 (figure 4I and table 4). For the SOTs, however, I were unable to acquire transcript data, as it has proven difficult to design specific qRT-PCR primers for these three transcripts due to their sequence similarity. Despite their high similarity, I acquired protein abundance data providing insights into the protein levels of the three SOTs. The comparison between transcript and protein expression levels of GLS biosynthetic genes across the different transcription factor mutants revealed altered transcript-to-protein relationships for some GLS biosynthetic enzymes, and suggests yet unknown layers of gene regulation of these pathways.

### 2.5 Induction of GLS biosynthesis by exogenous MeJa

The phytohormone MeJa induces plant defense responses including GLS accumulation (Alvarez et al., 2008; Schweizer et al., 2013; Goossens et al., 2016). I treated Arabidopsis seedlings with MeJa to study the dynamics of the GLS pathway enzymes upon induction. I observed no consistent changes in GLS levels or composition within the first twelve hours after induction, but thereafter, total GLS amounts steadily increased (figure 5A), suggesting that the response has a lag phase in which signaling, gene expression and metabolic rewiring occur. The observed increase in total GLS was entirely due to IG, which make up more than 80 % of total GLS after 24 hours (figure 5B). In fact, a slight decrease was observed in absolute levels of several AG upon induction with MeJa (supplemental figure S3 and supplemental table S4).

**Figure 5.**
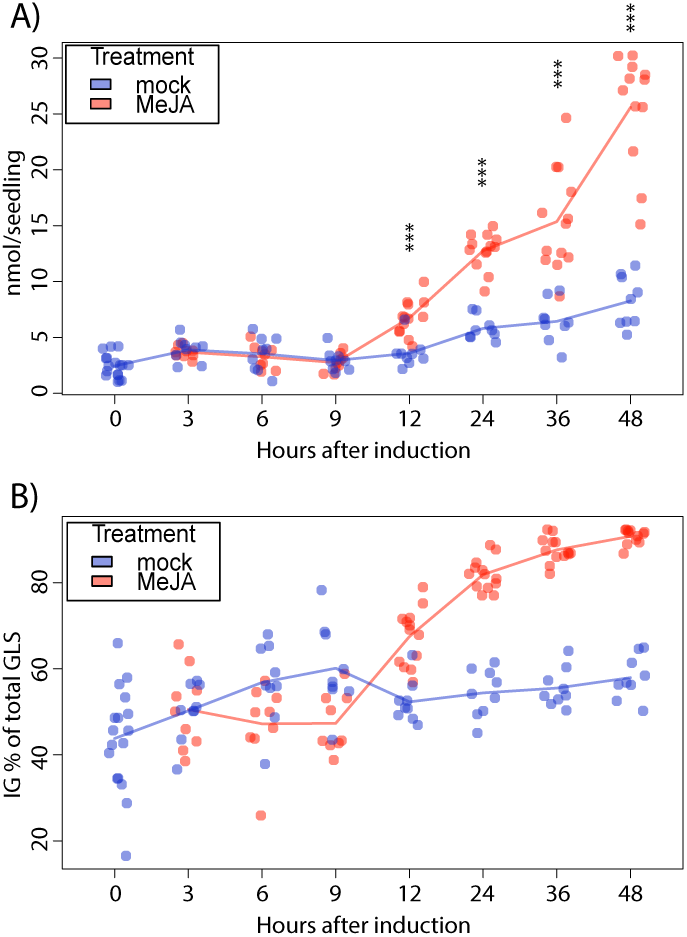
Induction of GLS accumulation by MeJa treatment. **(A)** Total GLS (nmol seedling^−1^) at different time points after MeJa treatment. **(B)** Percentage of IG of total GLS. Each dot represents a biological replicate (n = 10); lines represent means across these replicates. Asterisks denote statistical significant differences between mock and MeJa treatment at individual timepoints \*\*\**p*-value < 0.001, Welch two sample t-test as *posthoc* test, supplemental table S7)

To examine the events of the initial lag phase more closely, I extracted RNA and protein from MeJa-treated seedlings at different time points after induction. Upon MeJa treatment all measured transcripts responded with a rapid increase already one hour after treatment (figure 6 and table 5). Contrary to these observations, none of the corresponding enzymes was significantly increased upon treatment with MeJa (table 6). However, 25 hours after treatment, the two enzymes CYP83A1 and UGT74C1 showed a significant decrease in protein abundance (figure 6A, E and table 6). Our data reveal a discrepancy between the changes observed at the transcript and protein level in response to MeJa (figure 6 and table 6). While the transcripts responded fast and with high amplitude to MeJa treatment, protein levels remained largely unaffected.

**Table 5.**
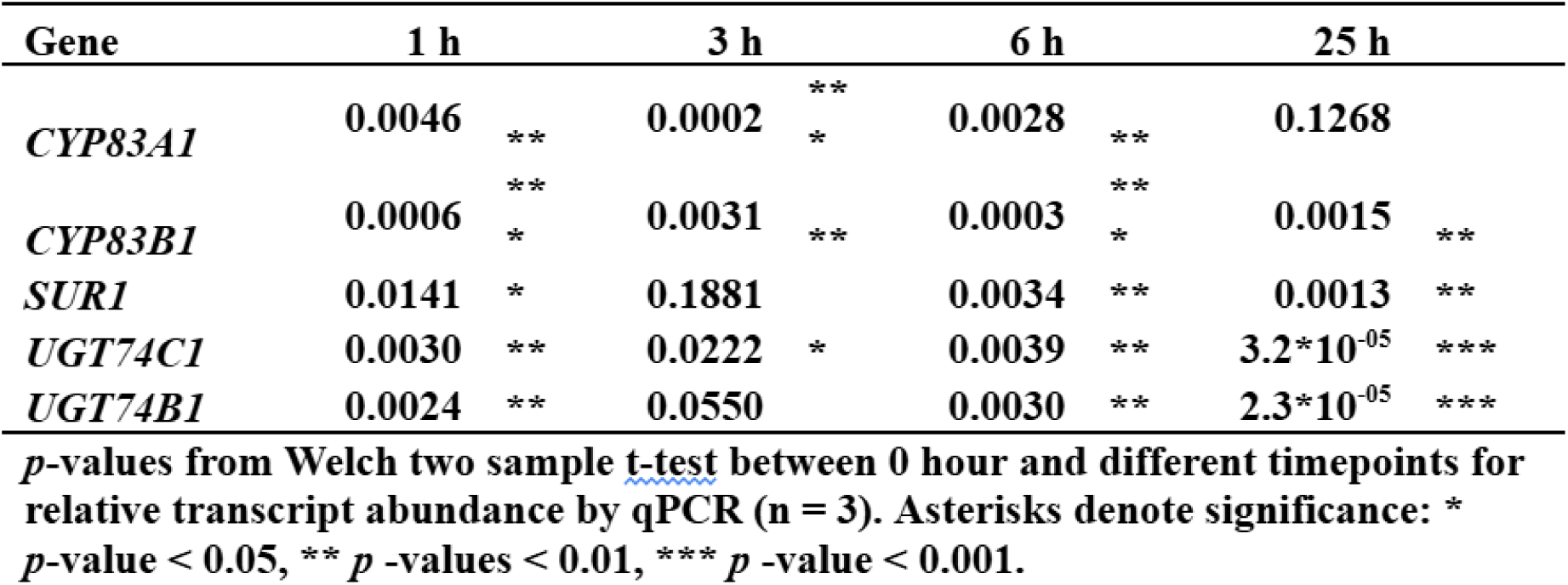
Statistical analysis of transcript changes upon MeJa treatment. Comparison betw een relative transcript abundance at 1,3, 6 and 25 hours after induction.

**Table 6.** Statistical analysis of protein changes upon MeJa treatment. Comparison between relative protein abundance at 1, 3, 6 and 25 hours after induction.

| **Protein** | **1 h** | **3 h** | **6 h** | **25 h** |  |
| --- | --- | --- | --- | --- | --- |
| **CYP83A1** | **0.1028** | **0.8799** | **0.164** | **0.0471** | **\*** |
| **CYP83B1** | **0.8075** | **0.4373** | **0.6048** | **0.9156** |  |
| **SUR1** | **0.3635** | **0.4591** | **0.7239** | **0.9039** |  |
| **UGT74C1** | **0.0271** | **0.6209** | **0.0662** | **0.0019** | **\*\*** |
| **UGT74B1** | **0.1739** | **0.9898** | **0.5381** | **0.1935** |  |

*p*-values from Welch two sample t-test between 0 hour and different timepoints for relative protein abundance by targeted proteomics (n = 3). Asteriks denote significance: \* *p*-value < 0.05, \*\* *p* -values < 0.01, \*\*\* *p* -value < 0.001.

**Figure 6.**
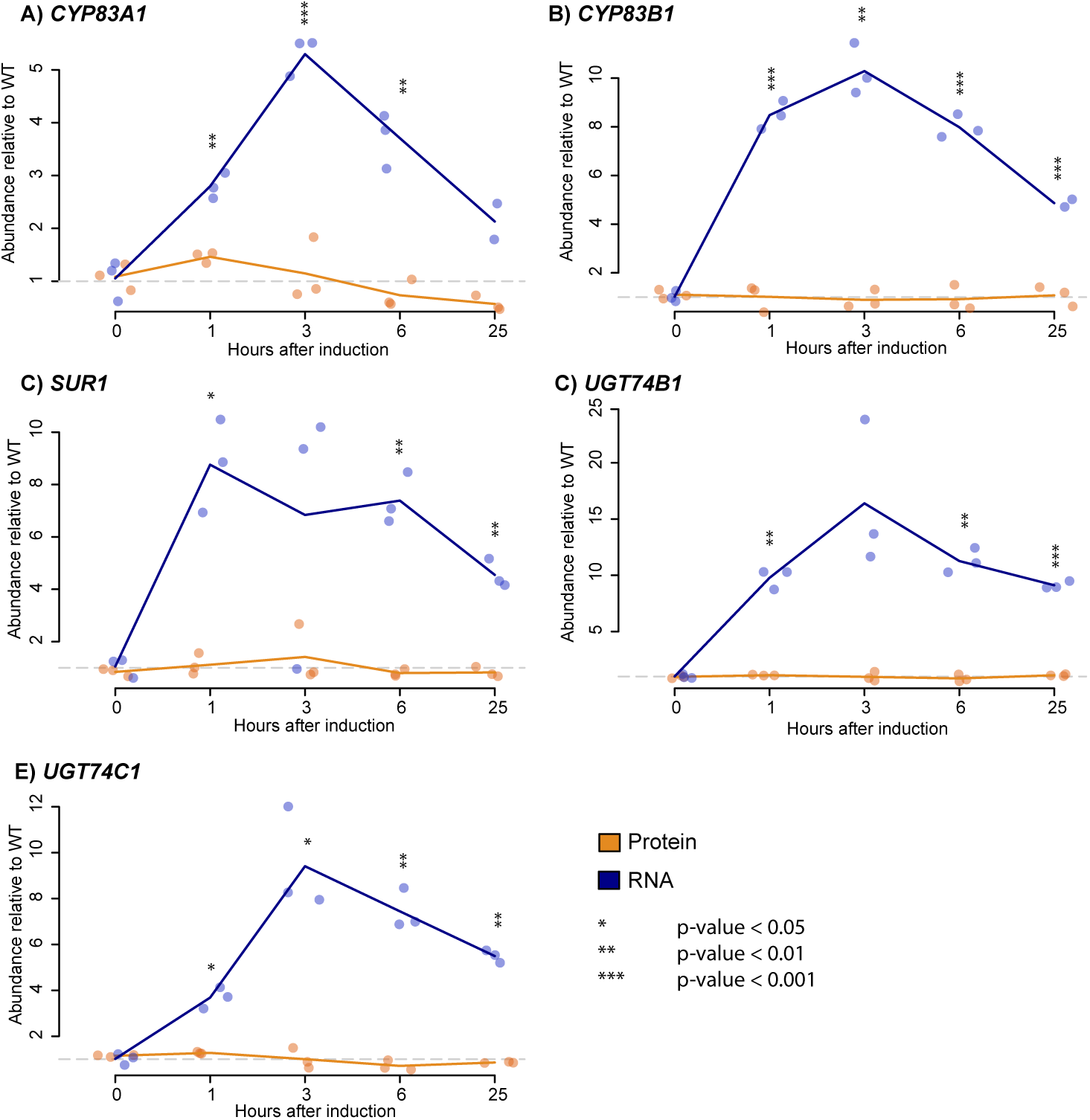
MeJa induced uncoupling of transcript and protein levels. Protein (yellow) and transcript (blue) were quantified at different time points after induction with MeJa for selected genes of GLS biosynthesis: (A) *CYP83A1*, (B) *CYP83B1*, (C) *SUR1*, (D) *UGT74B1* and (E) *UGT74C1*. Abundance is normalized relative to 0 hours after induction. Grey dashed lines represent relative abundance at 0 hours after induction. Asterisksdenote statistical significant differences in protein and RNA levels relative to time point 0 (\**p* -value < 0.05, \*\**p* -value < 0.01, \*\*\**p*-value < 0.001, Welch two sample t-test, n = 3, supplemental table S9).

## 3 Discussion

Using standard-flow driven targeted proteomics for the quantification of enzymes in GLS biosynthesis in Arabidopsis, I observed protein level changes across several transcription factor mutants impaired in GLS biosynthesis and upon induction of the pathways. Contrary to the large-scale DDA proteomics experiments – which require highresolution instrumentation such as Orbitrap instruments coupled to nano-flow LC – targeted proteomics requires a relatively simple and inexpensive triple quadrupole MS, which combined with standard-flow LC allows for reliable and versatile protein quantification.

### 3.1 Quantification of low-abundant cytochrome P450 enzymes

I successfully designed and implemented SRM assays covering enzymes of the core GLS pathways. With few exceptions, the peptides have a lower limit of quantification below 5 fmol pr. injection – very close to their detection limit. Even with the relatively high sensitivity of our assays it has been impossible to detect endogenous tryptic peptides of CYP79B2, CYP79B3, CYP79F2 and CYP71B15/PAD3. To our knowledge, these proteins have hitherto eluded proteomic detection by both shotgun and targeted proteomics. This could be explained by low expression levels, i.e. the peptides are below detection limit, which is well in line with the published transcript data (Schweizer et al., 2013; Burow et al., 2014). For biosynthesis of cyanogenic glucosides, a recent proteomics study in sorghum showed that the CYP79 enzyme catalyzing the first committed step in the pathway is far less abundant than the subsequent cytochrome P450 (Laursen et al., 2016). These observations support the idea of a ‘gatekeeper’ role for the initial cytochrome P450 enzymes, through which the pathway flux is restricted and controlled.

It is interesting that in our experiments neither genetic perturbations nor hormone treatment revealed the undetected cytochrome P450 enzymes. For CYP71B15/PAD3 a 20-fold increase in transcript levels has been reported in the myb34/51/122 mutant (Frerigmann et al., 2015), and MeJa treatments have been shown to severely increase the transcript levels of both CYP79B2 and CYP79B3 (Mikkelsen et al., 2003). As the cytochrome P450s are membrane-bound, it is possible that our difficulties detecting them arise from inefficient extraction and loss due to sample handling, as it can be a problem for proteomics of membrane-bound proteins. However, as three out of seven cytochrome P450s (i.e. CYP79F1, CYP83A1 and CYP83B1) are successfully detected in our assays, it seems unlikely to be due to inherent problems of analyzing this protein family. Instead, some cytochrome P450s may be expressed at levels below our detection limit even under induced conditions, which suggests strict regulation of translation rates or rapid protein turnover – an idea that is supported by recent results showing that the GLS enzyme levels are controlled by the ubiquitin-26S proteasome system (Svozil et al., 2015). Another interpretation could be widespread post-translational modification of the enzymes. Post-translational modifications of cytochrome P450 enzymes are known to occur in humans, and also in plants (Aguiar et al., 2005; Hayashi and Harada, 2014; Northey et al., 2016). Chemical modifications change the mass-to-charge ratio of the peptide, giving rise to different fragmentation patterns and chromatographic behavior and would render the designed SRM assays inapplicable. However, representing a protein by several peptides – as done in the present study – would protect against this problem, unless the protein of interest is modified at each individual proteotypic peptide.

### 3.2 Expression levels in transcription factor mutants

For our protein-transcript comparison in the three transcription factor mutants, it is important to emphasize that the relative quantification applied to both qRT-PCR and SRM assays does not give absolute levels of either RNA or protein. Instead I observe the relative differences between wild-type and the mutants. Our results show that for most genes the reduced transcript levels in the mutants correlate with a corresponding reduction in protein abundance, suggesting a simple relationship between transcription, translation and protein degradation (figure 4). Notable exceptions are UGT74C1 and SUR1.

Our results suggest that regulation of UGT74C1 is predominantly achieved via modulation of protein, rather than transcript level. However, as UGT74C1 is only quantified using a single proteotypic peptide, care should be taken when concluding on its abundance. UGT74C1 transcript levels were hardly affected in any of the three mutants, indicating that this gene is not strictly associated with GLS biosynthesis. However, it has been suggested that UGT74C1 is more involved in AG than IG biosynthesis (Grubb et al., 2014). Our results support this idea as the protein level is reduced in myb28/29 and myc2/3/4, but not myb34/51/122 (figure 4F).

For SUR1, transcript and protein levels showed the same patterns in both myb28/29 and myb34/51/122. However, in the myc2/3/4 mutant, the SUR1 protein level was more severely reduced (figure 4E). Besides regulation of transcript abundance, SUR1 is subject to several other regulatory mechanisms, including a microRNA-mediated regulation (Kong et al., 2015) and alternative splicing (www.arabidopsis.org, [11/10/2017]). These known modes of regulation, however, do not explain the observed transcript-protein discrepancy. microRNA targeting leads to reducedtranscript levels and the SRM assays are sensitive to alternative splicing, as the EENLVFLPGDAL-GLK peptide is unique to one splice variant, while the remaining two peptides IGWIALNDPEGVFETTK and FASIVPVLTLAGISK are present in both splice variants. I observed no discrepancy between EENLVFLPGDAL-GLK and the two other peptides (figure S3). Our results thus suggest myc2/3/4-dependent regulation of SUR1 specifically at the post-translational level.

Methyl-jasmonate induced uncoupling of transcript and protein levels in GLS biosynthesis Treatment with exogenous MeJa led to an increase in IG (figure 5), which was reflected at the transcript levels of the genes in IG biosynthesis (figure 6A). However, the genes in AG biosynthesis show similar increase in transcripts upon treatment, without translating into increased AG metabolite levels 25 hours after treatment. Quantification of the biosynthetic enzymes using SRM assays revealed no significant increases in IG enzyme levels in response to the treatment, whereas the AG enzymes decreased after 25 hours. This indicates that increasing the amount of enzymes is not required for increasing IG accumulation. A higher flux through the pathway, without a corresponding increase in enzyme abundance can be achieved by stimulating catalytic activity, modulating substrate availability or alleviating bottlenecks. To our knowledge, post-translational modification resulting in increased or decreased catalytic activity of any of these enzymes has not been reported; however, phosphorylation at several sites has been reported for UGT74B1, opening up for the possibility of post-translational regulation of this enzyme (Reiland et al., 2011; Wang et al., 2013).

Alternatively, as several catalytic steps are shared between the two pathways, a common metabolic bottleneck may exist due to scarcity of one or more of the shared substrates (glutathione, uracil-diphosphate glucose and 3’-phosphoadenosine-5’-phosphosulfate). Reducing the flux through one of the pathways may alleviate this bottleneck and allow for a faster flux in the other pathway. This aligns with our observation that treatment with MeJa led to a decrease in the protein levels of specifically CYP83A1 and UGT74C1 (enzymes associated with AG biosynthesis) after 25 hours of treatment (figure 6A and E), as well as the decrease in individual AG upon induction (supplemental figure S3 and supplemental table S4).

Reciprocal negative crosstalk between the two GLS pathways in Arabidopsis has been observed and discussed previously (Grubb and Abel, 2006; Malitsky et al., 2008). As described above, MeJa treatment led to a suppression of AG-specific enzymes. Similarly, increased levels of IG have been observed in cases with reduced AG-specific gene expression as in the myb28/29, ref2-3 and myb34/51/122 mutants (Kliebenstein et al., 2005; Sønderby et al., 2010a; Frerigmann and Gigolashvili, 2014). The exact nature of this reciprocal crosstalk remain unknown, but as electrons are required to drive the catalytic reactions of the cytochrome P450s, it has been hypothesized that this is the rate-limiting step of GLS biosynthesis (Grubb and Abel, 2006).

Support for the “limiting electron hypothesis” comes from the enigmatic cytochrome b5 proteins, which are known to – among other things – support cytochrome P450s with electrons. Higher plants have several cytochrome b5 proteins, and it is possible that these evolved to support the vast number of cytochrome P450s of higher plants. Recently, I were able to show that the cytochrome b5 isoform C of Arabidopsis affects particularly the accumulation of long-chained AG upon prolonged treatment of MeJa (Vik et al., 2016). It is possible that the cytochrome b5 proteins are important sources for electrons and that mutation of them disturb the catalytic activity of cytochrome P450s. Accordingly, down-regulating CYP83A1 enzyme levels would allow more of these electrons to be used by the cytochrome P450s involved in IG biosynthesis. However, as AG- and IG-specific enzymes show overlapping but not identical protein expression patterns (Nintemann et al., 2018), direct metabolic crosstalk can only occur in cells harboring both pathways.

Our experiments show that the large changes observed in transcript in response to MeJa treatment are not representative of the protein level, stressing the need for the targeted proteomics methodology to measure protein abundance in a reliable way. However, it is important to note that the transcript levels appear to reflect the observed changes in metabolite levels much more closely, suggesting that more knowledge about protein level regulation is required to understand the dynamics of GLS pathway orchestration.

## 4 Conclusion

Targeted proteomics allows investigation of specific proteins in different genetic backgrounds and under various conditions. Reliable quantification and ease of multiplexing has made targeted proteomics a central methodology for pathway engineering in synthetic biology (Redding-Johanson et al., 2011; Batth et al., 2014; González Fernández-Niño et al., 2015). In the presented work, quantification of enzymes in the GLS biosynthetic pathways provided valuable information about changes in enzyme levels under genetic and hormonal perturbations. Our results illustrate the need to obtain protein level data to better understand the regulatory processes underlying the dynamics of pathway orchestration and raise questions regarding yet unknown regulatory steps in transcript and protein expression. As biological research is moving towards quantitative biology, I expect that targeted proteomics – with its ability to probe, detect and quantify a specific subset of the proteome – will play an increasing role and will contribute to advancing our understanding of quantitative relationships and dynamics in biology.

## 5 Methods

### 5.1 Peptide design and method development

Peptides were designed using the pep2pro server, and Arabidopsis Proteotypic Peptide Predictor (Baerenfaller et al., 2011; Hirsch-Hoffmann et al., 2012; Taylor et al., 2014). Synthetic peptides (so-called SpikeTides L) were acquired from JPT (Berlin) with iodoacetamide alkylation of cysteins and 13C15N isotopically labeled arginine and lysine at the C-terminal. Skyline 3.6 was used for assay development and later data analysis (MacLean et al., 2010). A spectral library for the Arabidopsis proteome was acquired from the Global Proteome Machine (uploaded 5/30/13) (Craig et al., 2004). Transitions for collision energy optimization were picked by their ranking in the spectral library when available. In cases where a spectral library was not available, the five largest fragments were picked. After collision energy optimization, the top three transitions were picked for the final assay. For initial optimization of source settings and chromatography, commercially available synthetic peptides were used (MSRT-1VL, Sigma-Aldrich).

### 5.2 Plant material and growth conditions

The following Arabidopsis genotypes was used WT (Col-0), myb28/29 (Sønderby et al., 2007), myb34/51/122 (Frerigmann and Gigolashvili, 2014) and myc2/3/4 (Schweizer et al., 2013). Seeds were sterilized by incubating in 70 % (v/v) ethanol, 0.05 % (v/v) Triton X-100 for 5 min and washing in 70 % (v/v) ethanol. The air-dried seeds were plated on ½ MS (Murashige and Skoog medium including vitamins; M0222, Duchefa) agar plates and cold-stratified for 48 hours at 4 °C in the dark. For the mutant analyses, seedlings were grown for 14 days at 20 °C under 16 hours of light with an intensity of 110-140 µE. Treatment with MeJa was done by carefully transferring 7 day old seedlings to ½ MS agar plates containing 200 µM MeJa (392707-5ML, Sigma-Aldrich). For each biological replicate, 50 mg of seedlings were collected in a 1.5-ml reaction tube containing two chrome balls and flash frozen in liquid nitrogen (material was stored at -80 °C until further processing). Each biological replicate was grown on an independent agar plate.

### 5.3 RNA and protein extraction and qRT-PCR

Plant tissue (50 mg) was ground to powder using a mixer mill and then processed with the RNeasy Plant Mini Kit (74904, Qiagen) according to manufacturer’s instructions. However, the flowthrough (containing protein) of the RNAspin columns was collected, acetone-precipitated according to manufacturer’s instructions (Qiagen, Supplementary protocol, RY22 Jan-06), and processed further described below: in solution protein digest. The extracted RNA was DNase treated using DNase 1 (AMPD1-1KT, Sigma-Aldrich) and reverse transcribed using the iScript cDNA synthesis kit (#170-8891, Bio-Rad). qRT-PCR was performed using the SYBR green dye (DyNAmo Flash SYBR green qPCR kit, F415S, Thermo Scientific) and the following primers:

CYP79F1

5’-TGTCCCTTCCCATCTTGCGCGT-3’ and
5’-ACGACCTAGTCCAGGGCGGC-3’;

CYP83A1

5’-TGTCATGACTGGTCTTGCTATGCAC-3’ and

5’-CATTGAAACAGAATACACTGGAGGAACA-3’;

CYP83B1

5’-GGAGTCTACCTAAAGGGATTAAACCAGAG-3’ and
5’-CATATCTACCAGCAGAAACGTCCTAATG-3’;

GGP1

5’-AGCTCTCACGATGCCTTTGAGAATG-3’ and
5’-ACCCTGGCTATGATCTGATGACCAA-3’;

SUR1

5’-GCACTTGAGAGACTGAAGGGTTTCTG-3’ and
5’-GAATATTCTTTTGGGCACACACATCC-3’;

UGT74B1

5’-GTGTGCCTCAGTGGAGTGAT-3’ and
5’-ACGATTACTTCCCCAGCTTCC-3’;

UGT74C1

5’-GTGGCTAAGTGGGTTCCTCA-3’ and
5’-CCCCTAAGCATAGTGCCTCC-3’.

PEX4/UBC22 (AT5G25760)

5’-CTGAGCCGGACAGTCCTCTTAACTG-3’ and
5’-CGGCGAGGCGTGTATACATTTGTG-3’ was used as reference (Czechowski et al., 2005). Relative quantification of changes in steady-state mRNA levels was done as previously described (Pfaffl, 2004).

### 5.4 In solution protein digest

The acetone precipitate was spun down at 3000 g for 20 min at 4 °C, after which pellets were washed twice with ice-cold acetone, 10 % methanol, 10 mM dithiothreitol. The resulting pellets were resuspended in 100 µL 8 M urea, 100 mM TrisHCl, pH 8.2. Protein content were quantified using the 660 nm protein assay (22660, Pierce) and 100 µg of total protein were collected in new 1.5-ml reaction tubes. To each sample, 2 µL of 10 mM dithiothreitol were added and the samples were gently shaken for 30 min at room temperature. 10 µL 55 mM iodoacetamide were added to the samples and they were incubated for 20 min in the dark. The samples were then diluted wiht 50 mM ammonium bicarbonate (typically 200-300 µL), to arrive at a urea concentration of 1.8 M. 2 µg of trypsin/lys C mix (V5073, Promega) were added to each tube, and the samples were incubated at 37 °C overnight. Digest was stopped by acidification by addition of 10 % triflouracetic acid until pH ¡ 2.5 (typically 10-20 µL). Samples were then diluted up to a final volume of 1 mL in 2 % acetonitrile, 0.1 % formic acid and spun at 20.000 g for 15 min. For each sample a sep-pak C18 cartridge (WAT023590, Waters) was prepared by washing with 1 mL of 70 % acetonitrile, 0.1 % formic acid, followed by 1 mL of 2 % acetonitrile, 0.1% formic acid. Peptide samples were applied followed by three washings with 2 % acetonitrile, 0.1% formic acid. The samples were finally eluted by two applications of 500 µL 70 % acetonitrile, 0.1 % formic acid, after which the samples were dried using a speedvac at 1000 rpm, 30 °C. The dried peptide pellets were stored at –20 °C until analysis. Prior to analysis, the pellets were resuspended in 25 µL 2 % acetonitrile, 0.5 % formic acid, 0.1 % triflouracetic acid containing synthetic, heavy labeled peptides at a concentration of 10 fmol/ µL. The samples were filtered through 0.22 µm spinfilters (516-0234, VWR) which were washed with additional 10 µL of 2 % acetonitrile, 0.5

% formic acid, 0.1 % triflouracetic acid.

### 5.5 Liquid chromatography and mass spectrometry for proteomics

The gradient was adopted from Percy et al. and Batth et al. with slight modifications (Percy et al., 2012; Batth et al., 2014). I utilized a gradient of buffer A (0.1 % formic acid) and B (99.9 % acetonitrile and 0.1 % formic acid). At a flow of 500 uL/min the percentage of B was as follows: 5 % at start; increased to 10% at 1 min; 11 % at 3 min; 19 % at 13 min; 27.5 % at 21 min; 34 % at 21.7min; 42 % at 22.5 min; 90 % at 23.50 min; kept at 90 % at 26.90 min; reduced to 5 % at 30 min; kept at 5 % until 34 min. Column oven was set to 55 °C. Peptide separation was achieved on an aeris PEPTIDE, XB-C18 column (1.7 µM particle size, 2.1 mm i.d. x 150 mm length, Phenomenex) on an Advance UHPLC-OLE (Bruker). The injection volume was 10 µL. Source settings for heated electrospray ionization were as follows: spray voltage 3200 V, positive mode; cone temperature 300 °C; cone gas flow 20 psi; heated probe temperature 300 °C; probe gas flow 40 and nebulizer gas flow 50. The triple quadrupole mass spectrometer (EVOQ Elite, Bruker) was set to scan for transitions for individual peptides within scheduled 2 min windows. Resolution of the first and third quadrupole was set to 1 Da. The individual retention times and transitions are found in table 1. The acquired chromatograms were manually inspected and peak area ratios between endogenous light and synthetic heavy peptides were obtained through Skyline 3.6. Undetected peaks (N/A) were treated as zero values for analysis purposes. Additional statistical analysis and visualization was performed in R studio v0.99.892 (RStudio Team, 2015).

### 5.6 Glucosinolate extraction and analysis

Seedlings were harvested for GLS extraction by submerging single seedlings in tubes with 300 µL 85 % methanol containing 1 nmol p-hydroxybenzyl GLS as internal standard and a chrome ball. Tissue was ground using mixer mill and supernatants were collected using an anion exchange resin (DEAE Sephadex, A-25, GE Healthcare). Eluates obtained by sulfatase treatment were diluted 1:10 and analyzed by LC-triple quadrupole MS as described previously (Crocoll et al., 2016).

## 6 Acknowledgements

I would like to thank Barbara A. Halkier and Meike Burow for supporting me in the thesis work which led to this paper; Henriette S. K. Jepsen for excellent technical assistance with qRT-PCR; Sebastian J. Nintemann for providing CYP83B1 primers for qRT-PCR; Tamara Gigolashvili and Philippe Reymond for providing seeds of myb34/51/122 and myc2/3/4, respectively. Financial support for the work was provided by the Danish National Research Foundation DNRF grant 99.

---

**Figure S1.**
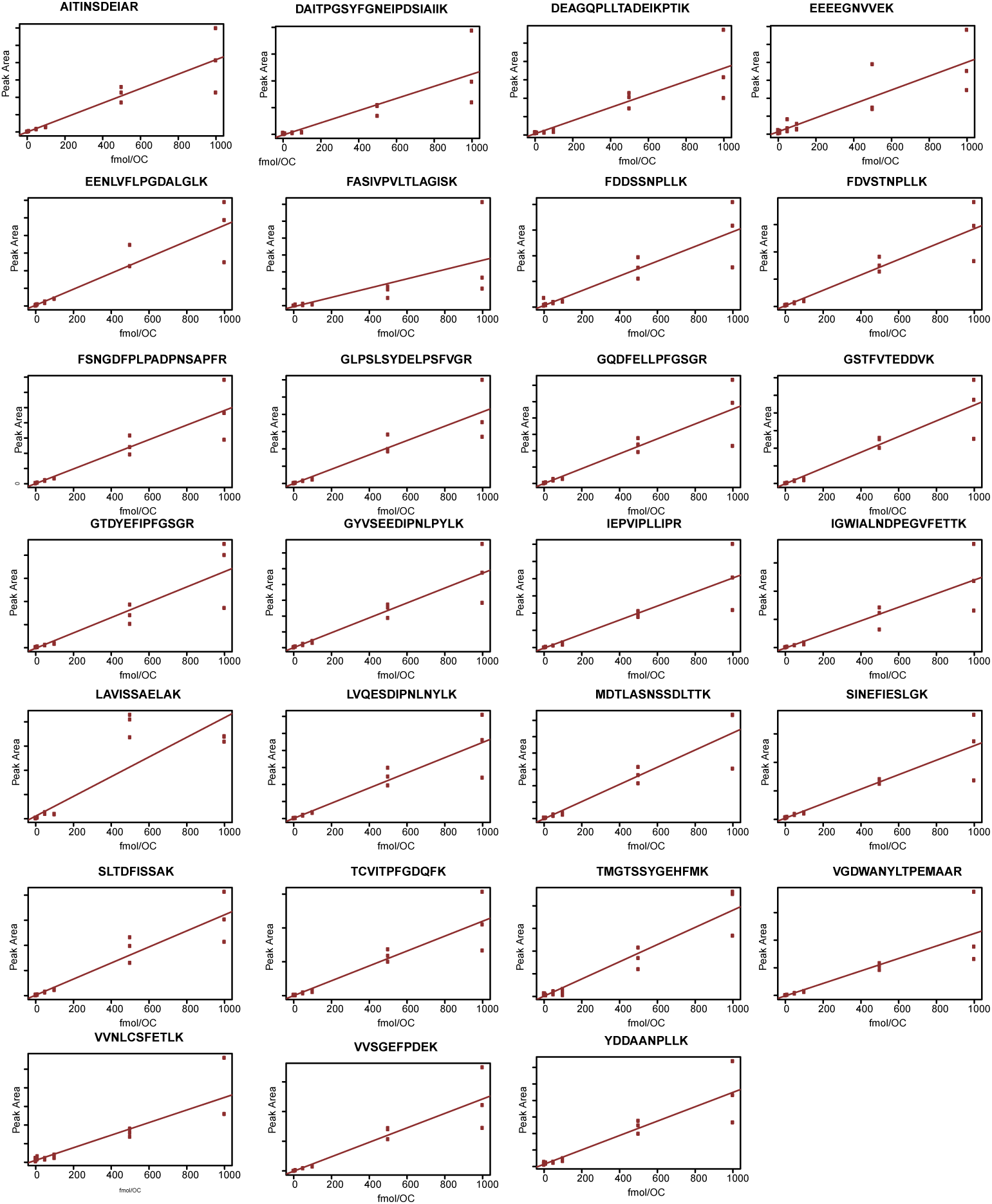
Standard curves for SRM peptides. Protein extracts from Arabidopsis seedlings spiked with varying concentrations of stable-isotope lablled synthetic peptides. Peak area as function of concentration (fmol on column). Solid line are the linear regression fitted to the datapoints.

**Figure S2.**
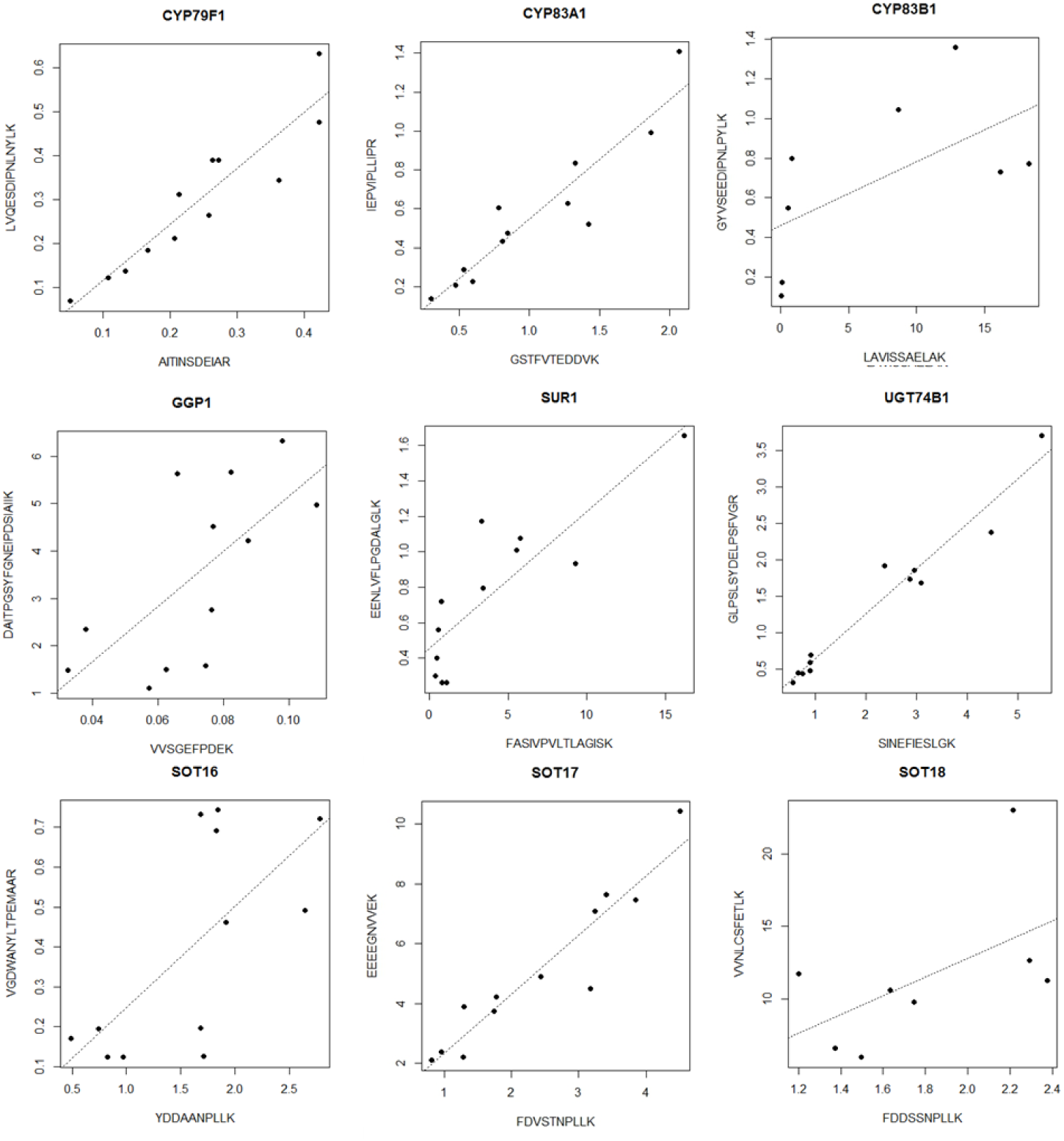
Peptide-peptide plots. The light-to-heavy ratios for two peptides are plotted against each other. Dashed line represent the linear regression.

**Figure S3.**
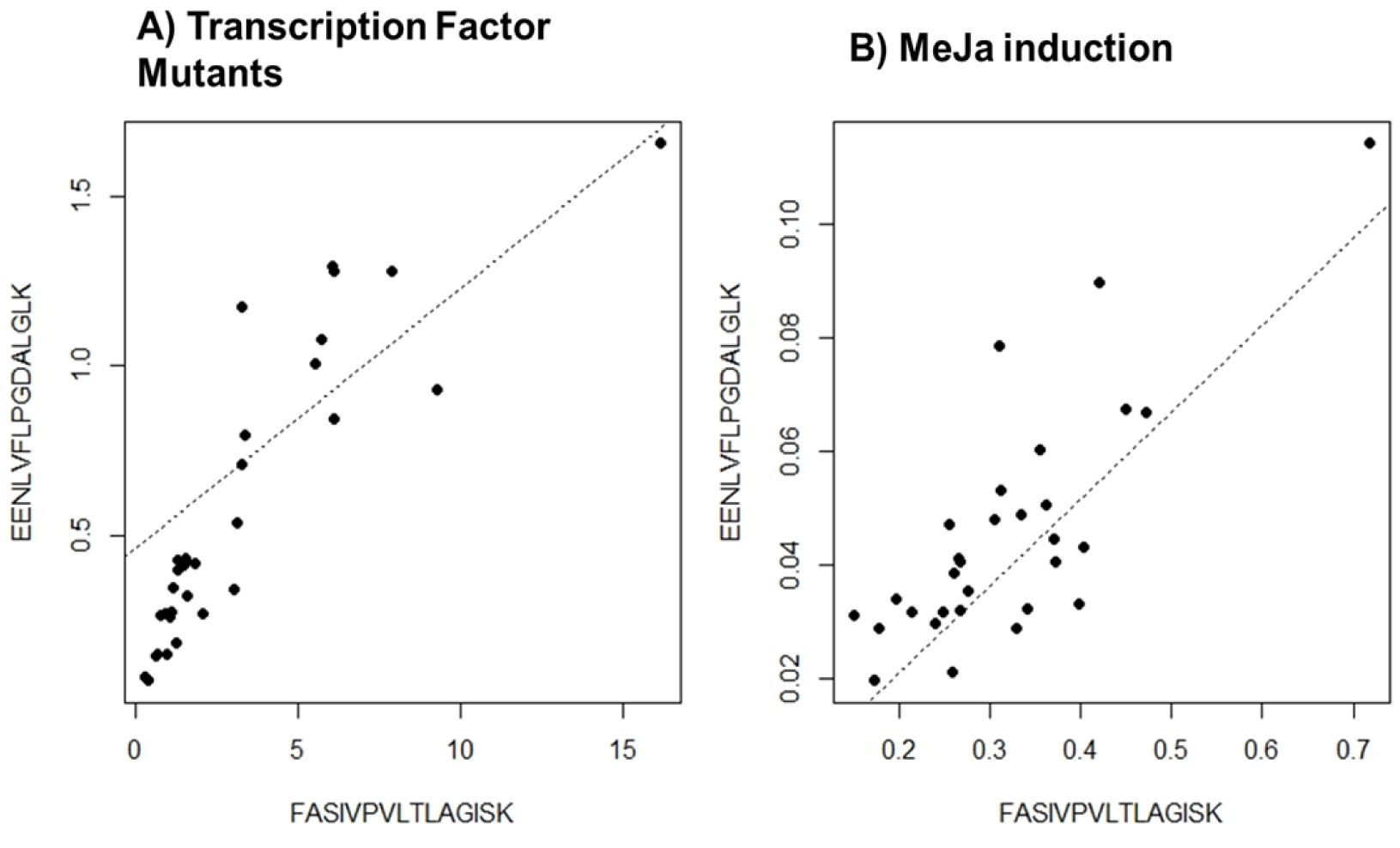
Peptide-peptide plots. The light-to-heavy ratios for two peptides of SUR1 are plotted against each other. Dashed line represent the linear regression. A) Is the data from transcription factor mutants. B) is the data from MeJa induction.

**Table S4.** ANOVA table for statistical analysis of transcript changes in mutants. p-values reported in ANOVA(RNA level = genotype + exp. + exp.:genotype)

| gene | genotype | exp. | exp.:genotype |
| --- | --- | --- | --- |
| CYP79F1 | 1.03E-11 *** | 0.5173 | 0.0310 * |
| CYP83A1 | 6.69E-18 *** | 0.0309 * | 0.0771 * |
| CYP83B1 | 4.53E-11 *** | 0.5412 | 0.9666 |
| GGP1 | 5.68E-07 *** | 0.9059 | 0.9709 |
| SUR1 | 1.66E-02 * | 0.7663 | 0.8496 |
| UGT74C1 | 6.30E-04 *** | 0.1908 | 0.7908 |
| UGT74B1 | 2.03E-09 *** | 0.6494 | 0.6912 |

\*p-value < 0.05, \*\* p-value < 0.01, \*\*\* p-value < 0.001

**Table S5.**
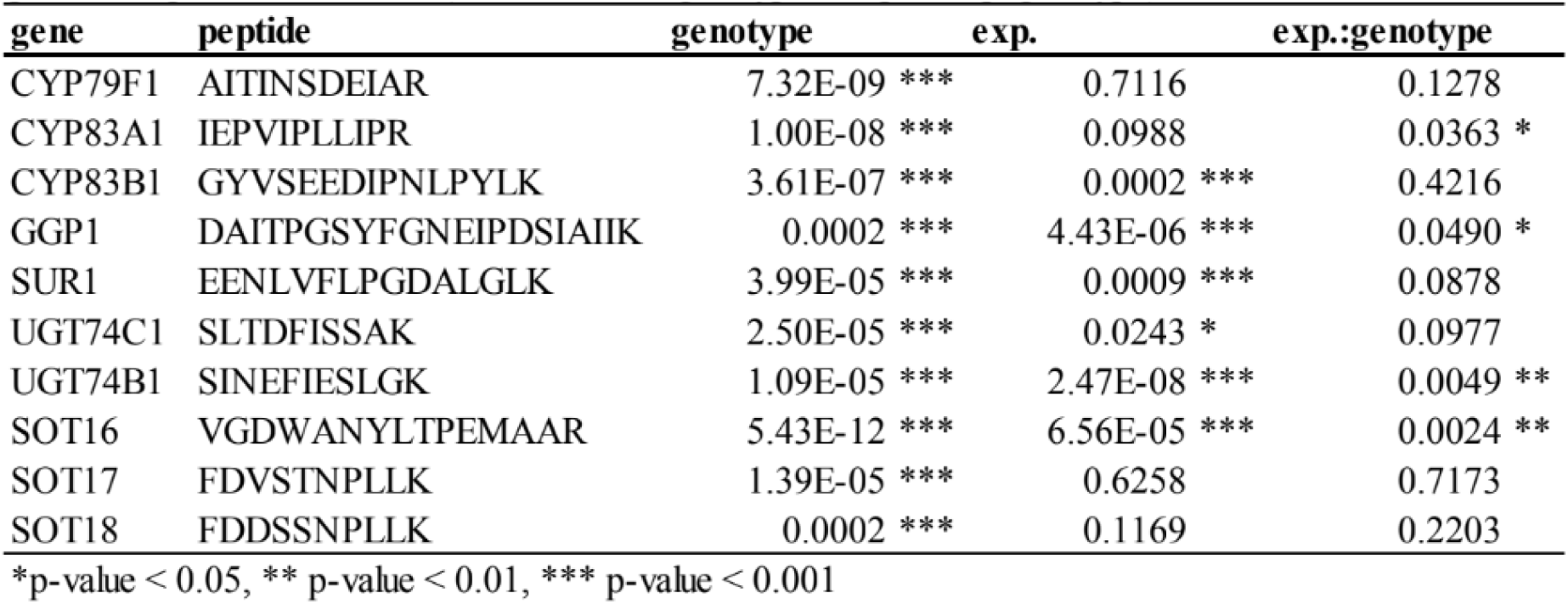
ANOVA table for statistical analysis for protein changes mutants. p-values reported in ANOVA(Protein level = genotype + exp. + exp.:gcnotypc)

**Table S6.**
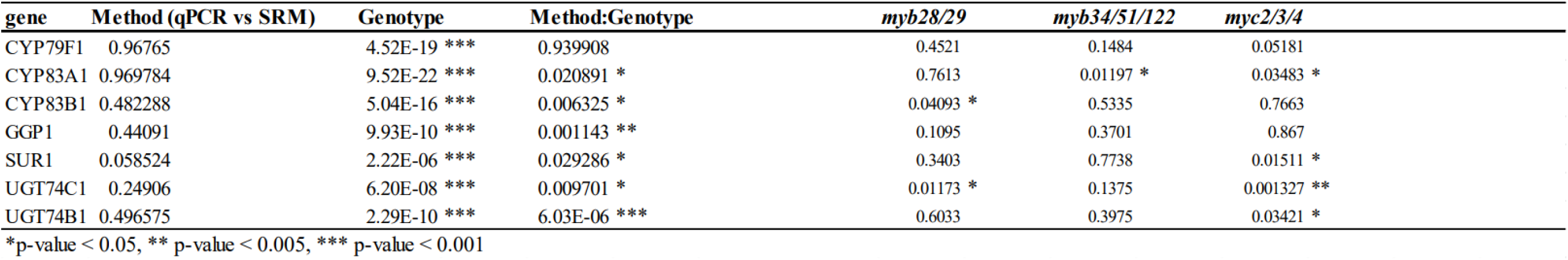
Statistical analysis of discrepancy between transcript and protein levels across genotypes. p-values reported in ANOVA (expression level = method + genotype + method:genotype) p-values from *posthoc* test (Method:Genotype, Welch two sample t-test)

**Table S7.** Statistical analysis of changes in individual glucosinolates upon MeJa treatment Hours after induction.

| GLS | 3 | 6 | 9 | 12 | 24 | 36 | 48 |
| --- | --- | --- | --- | --- | --- | --- | --- |
| ds.3mtp.I | 0.867 | 0.621 | 0.290 | 0.015 | 0.409 | 0.717 | 0.030 |
| ds.3msp.I | 0.547 | 0.327 | 0.835 | 0.760 | 0.173 | 0.395 | 0.537 |
| ds.3ohp.I | 0.768 | 0.866 | 0.944 | 0.022 | 0.521 | 0.347 | 0.200 |
| ds.3bzo.H | 0.956 | 0.728 | 0.527 | 0.387 | 0.676 | 0.017 | 0.145 |
| ds.4mtb.I | 0.660 | 0.784 | 0.164 | 0.146 | 0.131 | 0.020 | 0.002 |
| ds.4msb.I | 0.278 | 0.857 | 0.695 | 0.422 | 0.935 | 0.099 | 0.008 |
| ds.4ohb.I | 0.132 | 0.952 | 0.206 | 0.511 | 0.804 | 0.220 | 0.141 |
| ds.4bzo.H | 0.885 | 0.089 | 0.592 | 0.602 | 0.903 | 0.061 | 0.196 |
| ds.5msp.I | 0.292 | 0.964 | 0.956 | 0.693 | 0.470 | 0.035 | 0.003 |
| ds.7mth.I | 0.305 | 0.047 | 0.092 | 0.033 | 0.251 | 0.013 | 0.731 |
| ds.7msh.I | 0.299 | 0.399 | 0.866 | 0.316 | 0.756 | 0.018 | 0.125 |
| ds.8mto.I | 0.014 | 0.099 | 0.011 | 0.158 | 0.186 | 0.377 | 0.182 |
| ds.8mso.I | 0.465 | 0.925 | 0.813 | 0.411 | 0.204 | 0.086 | 0.412 |
| ds.I3M.H | 0.068 | 0.011 | 0.001 | 2.48E-07 | 4.00E-10 | 2.58E-06 | 1.84E-06 |
| ds.4MOI | 0.952 | 0.809 | 0.471 | 0.371 | 0.338 | 0.013 | 0.118 |
| ds.NMOI | 0.954 | 0.634 | 0.304 | 1.29E-05 | 4.44E-11 | 3.33E-07 | 5.20E-09 |

*p*-values from Welch two sample t-test comparing GLS levels in mock and MeJA treated seedlings at each timepoint. Red colour highlight *p*-values below 0.05

**Figure S8.**
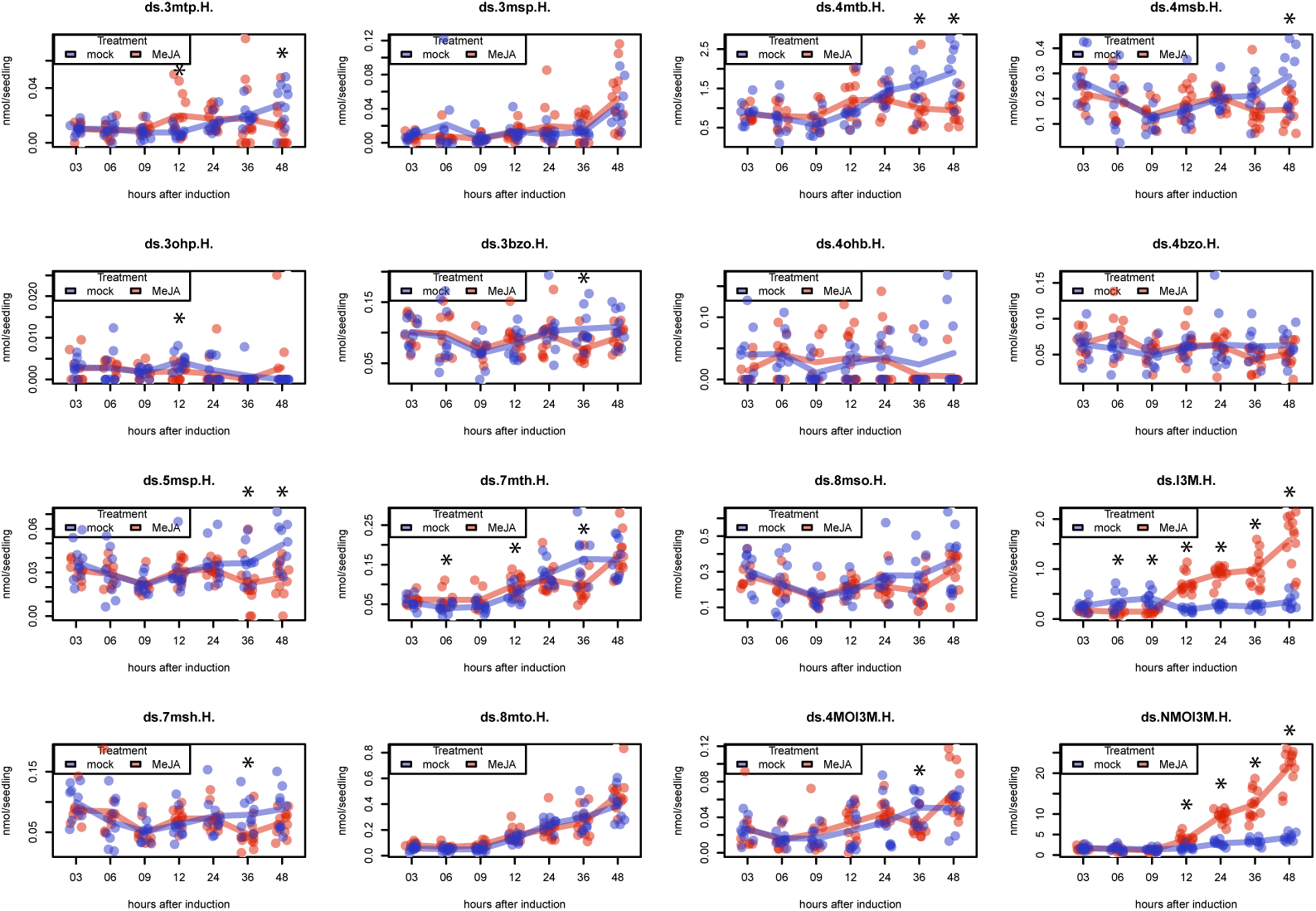
Changes in individual GLS in response to MeJa treatment. Arabidopsis seedlings treated with MeJa and subsequently analyzed for GLS content at different timepoints. Dots represent biological replicates, lines represent the mean levels. Colours blue and red correspond to mockor MeJa treated seedlings, respectively. Asteriks denote statistical significance by Welch two sample t-test (\**p*-value < 0.05). Details of analysis can be found in supplementary table S7.

**Table S9.**
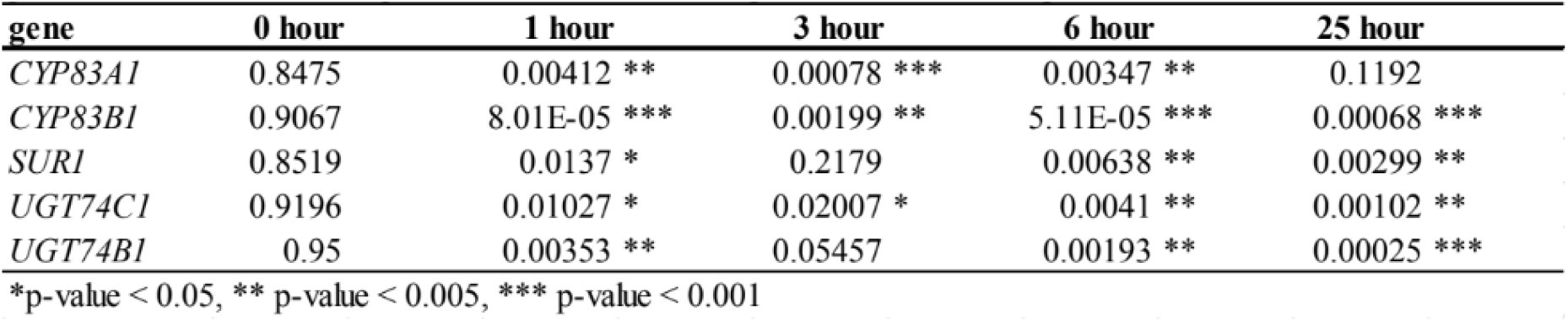
MeJa-induced discrepancy between RNA and protein. p-values from welch two sample t-test between RNA and protein at different timepoints

| **gene** | **0 hour** | **1 hour** | **3 hour** | **6 hour** | **25 hour** |
| --- | --- | --- | --- | --- | --- |
| *CYP83A1* | 0.8475 | 0.00412 ** | 0.00078 *** | 0.00347 ** | 0.1192 |
| *CYP83B1* | 0.9067 | 8.01E-05 *** | 0.00199 ** | 5.11E-05 *** | 0.00068 *** |
| *SUR1* | 0.8519 | 0.0137 * | 0.2179 | 0.00638 ** | 0.00299 ** |
| *UGT74C1* | 0.9196 | 0.01027 * | 0.02007 * | 0.0041 ** | 0.00102 ** |
| *UGT74B1* | 0.95 | 0.00353 ** | 0.05457 | 0.00193 ** | 0.00025 *** |

\*p-value < 0.05, \*\* p-value < 0.005, \*\*\* p-value < 0.001

